# Advanced Extracellular Vesicle Isolation: A Hybrid Electrokinetic-Tangential Flow Filtration Approach for Improved Yield, Purity, and Scalability

**DOI:** 10.1101/2025.03.27.645827

**Authors:** YongWoo Kim, SoYoung Jeon, KangMin Lee, Sehyun Shin

## Abstract

As extracellular vesicles (EVs) become increasingly important in diagnostics and therapeutics, achieving both high purity and yield during isolation remains a critical challenge. Conventional techniques often suffer from the co-isolation of non-vesicular particles and soluble proteins, limiting their clinical and research utility. In response, we introduce ExoTFF, a hybrid isolation technology that sequentially integrates electrokinetic filtration (ExoFilter) with size-exclusion tangential flow filtration (TFF) to deliver unprecedented performance gains through an iterative, synergistic mechanism. In the ExoTFF system, the sample is repeatedly circulated through an electrokinetic mesh filter and TFF until the liquid is removed. This recirculating flow gradually eliminates contaminants, while the electrokinetic filter continuously captures EVs as the sample is purified. Finally, any residual impurities in the TFF unit are completely removed via a dead volume elimination process. The complementary actions of these two distinct separation mechanisms double EV recovery rates and reduce impurity levels by 80% compared to conventional TFF, culminating in an impressive 800% improvement in the purity ratio. In proof-of-concept experiments, ExoTFF processed 10 mL of plasma within 10 minutes, efficiently depleting albumin and HDL while achieving superior EV recovery. To further explore scalability, an automated ExoTFF system processed 500 mL of sample in 50 minutes, maintaining consistent yield and purity. The ability to sustain performance across different scales highlights ExoTFF’s potential for both laboratory research and industrial-level EV production. Beyond biological applications, this platform also offers broad applicability for the isolation of negatively charged nanoparticles, demonstrating its potential impact across multiple nanotechnology-driven fields.

## 1. INTRODUCTION

Extracellular vesicles (EVs), the nanoscale particles enveloped by a lipid bilayer, are secreted by all types of cells and have emerged as vital mediators of intercellular communication.^1, 2^ Their ability to traverse biological barriers facilitates their roles in a variety of physiological and pathological processes, such as immune regulation, angiogenesis, and cancer metastasis.^3, 4^ Although EVs are small, they possess intrinsic properties that allow them to carry an array of biomolecules—including cytoplasmic and membrane proteins, lipids, and RNA molecules—which gives them tremendous potential for various applications, including targeted therapeutic delivery.^5^ Particularly, their natural capability to convey bioactive molecules position EVs as promising cargos for delivering drugs specifically to diseased cells.^3, 6^

As advancements in utilizing EVs for clinical and industrial applications continue, the need for high-purity purification technologies, along with large-scale isolation techniques, has become more crucial than ever.^7^ Among current purification methods, the combination of density gradient-based ultracentrifugation (UC) and size-exclusion chromatography (SEC) is the most commonly used approach for achieving high purity in EV isolation.^8–10^ However, these technologies are designed for small-scale laboratory samples and therefore exhibit significant limitations when scaled up for industrial applications.

While ultrafiltration and Tangential Flow Filtration (TFF) are among the technologies adopted for large-scale sample extraction in industry,^11^ these methods often fail to meet the high purity standards required in clinical settings.^12, 13^ Since each technique has its own advantages and limitations, and operates based on different isolation principles, a trend has emerged in recent publications to combine methods such as SEC with ultrafiltration (UF).^14, 15^ However, since both SEC and UF rely on the same size-based separation mechanism, these combined technologies are limited in their ability to remove impurities of similar sizes, inevitably restricting improvements in purity. In fact, nanoparticles such as HDL, which are abundant in samples like plasma, have been reported to remain unremoved).^14, 15^ Considering this, it is clear that a single separation technique alone cannot achieve sufficient purification. Moreover, when designing a combination of two or more techniques for purification, it is advisable to incorporate different isolation mechanisms that complement each other’s weaknesses while enhancing their strengths.^14^

Hence, this study aims to develop an innovative EV isolation technology that achieves high-level purification while maintaining high throughput. To this end, we carefully selected two technologies with different isolation mechanisms from among various available methods and integrated them through an advanced unified design into a single hybrid system, ExoTFF. The ExoTFF system consists of ExoFilter employing an electrokinetic-based separation mechanism and TFF utilizing a size exclusion principle. Recently, we reported ExoFilter, which is an electrokinetic filtration technology based on surface charge.^16^ When electrokinetic ExoFilter is combined with a size-exclusive TFF method, it produces an unexpectedly remarkable synergistic effect in isolation and purification. This synergistic effect arises from the integration of TFF’s impurity exclusion and liquid concentration capabilities with the ExoFilter. As TFF gradually removes impurities, the remaining sample becomes enriched with EVs, which are then more effectively captured by the ExoFilter. Moreover, the continuous recirculation of the sample through TFF increases the number of encounters between the target EVs and the ExoFilter. Consequently, the extraction efficiency is at least doubled compared to conventional TFF, impurity concentration is reduced to one-eighth, and the purity ratio is increased by 800%.

To apply this principle, this study developed an innovative syringe-type ExoFilter and connected it with a conventional cylindrical TFF system, thereby realizing a disposable ExoTFF system that integrates both technologies into one. To confirm the performance of the ExoTFF and its advantages over existing methods, we analyzed isolated EV on blood plasma samples according to ‘MISEV2023’,^17^ and compared the performance of ExoTFF with that of a single TFF or ExoFilter. We then extended the ExoTFF application to various biofluids, including urine, saliva, and culture medium. Additionally, as proof of concept for large-volume sample processing, an automated ExoTFF system was developed and demonstrated for handling 500 mL of plant extracted liquid within 10 minutes.

## 2. RESULTS

### Design of device and operating principle

This study proposed a hybrid technology that combines charge-based and size-based filtration methods to isolate high-purity extracellular vesicles (EVs), as illustrated in Figure 1(a). The charge-based filtration strategy, as shown in Figure 1(a-c), involves constructing a multi-layered, positively charged mesh, called ExoFilter, by binding cationic materials to the mesh.^16^ This ExoFilter selectively captures negatively charged EVs through electrokinetic filtration from other negatively charged proteins. Notably, the ExoFilter uses a mesh with 10 µm pores, allowing EVs to be captured without size exclusion, thereby minimizing mechanical stress associated with size-based filtration and preserving the structural integrity of the EVs.

**Figure 1.**
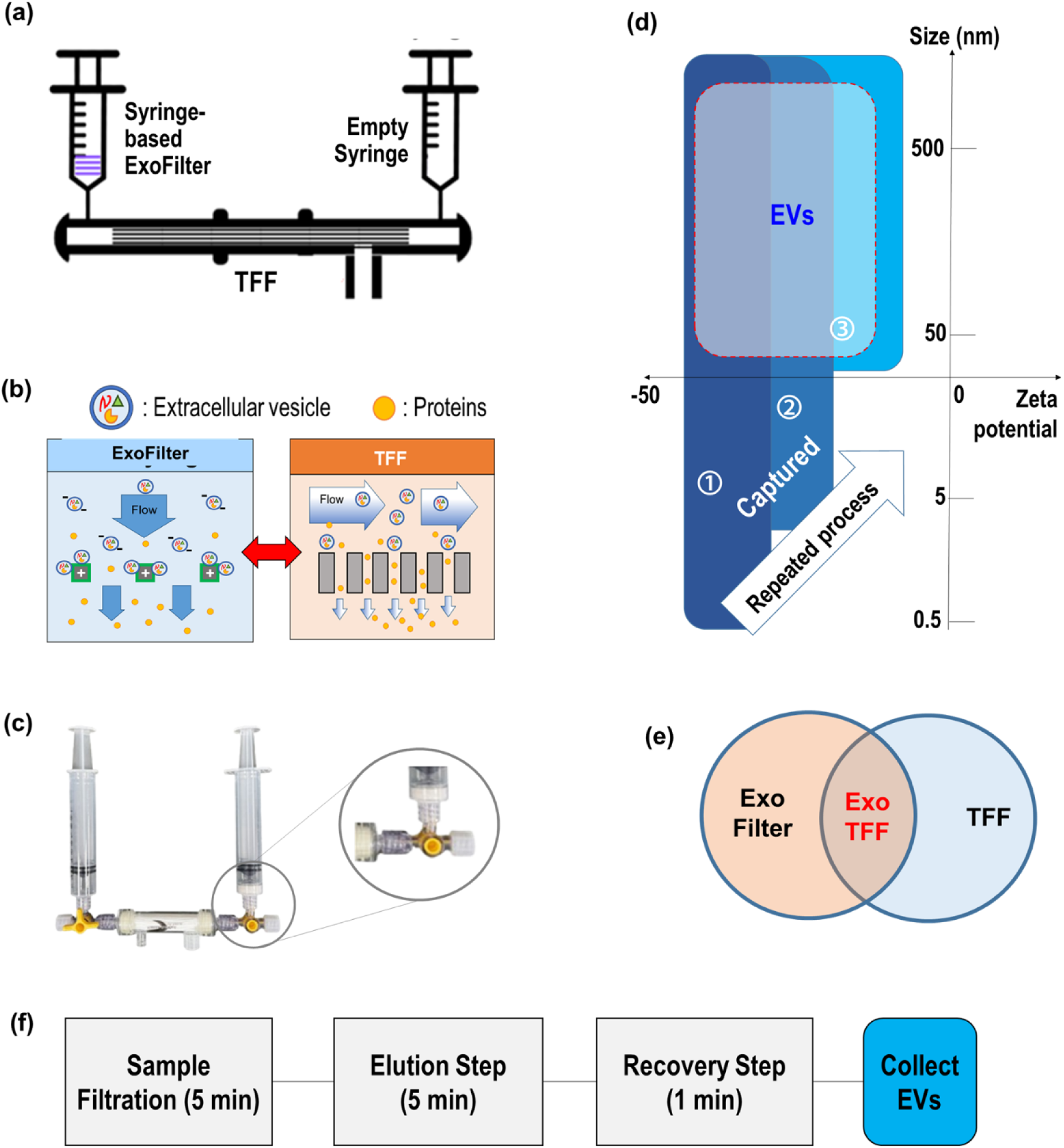
Hybridization of electrokinetic mesh filtration (ExoFilter) and size-based Tangential Flow Filtration (TFF). (a) Schematic of electrokinetic assisted mesh flow filtration with TFF, (b) Dual mechanisms of ExoTFF consisting electrokinetic filtration and size exclusion filtration, (c) A photographic image of the ExoTFF system, showing the syringe-based ExoFilter connected to TFF through a luer lock mechanism, (d) Operation window of ExoTFF in zetapotential vs. size domain, (e) Venn diagram of the hybrid ExoTFF system, which combines key features of ExoFilter and TFF, (f) Protocol of ExoTFF consisting of sample filtration, elution and recovery processes.

In the design of the present ExoTFF, we adopted size-based filtration, specifically TFF. As depicted in Figure 1(b), TFF uses hollow fibers with nanometer-sized pores to remove proteins smaller than the pore size while effectively isolating larger EVs. We used a commercial TFF (TFF-EVs, Hansa BioMed, Estonia) with an 800 kDa molecular weight (50 nm) cut-off. We explore the potential of a straightforward sequential process combining ExoFilter and TFF techniques, as shown in Figure 1s. For instance, we conducted two order-dependent experiments: one applying TFF followed by ExoFilter, and the other using ExoFilter followed by TFF. An interesting finding, detailed in the supplementary information (Fig. 1s), is that the sequence of the applications significantly affects both purity and yield.

Instead of operating the ExoFilter and TFF as separate sequential processes, we designed an innovative ExoTFF system that integrates two distinct systems—a syringe-based ExoFilter and TFF—into a single unified system, as shown in Figure 1(c). In fact, in conventional TFF operation, two syringes are connected to both ends of the TFF to drive the fluid manually. When one of these syringes is replaced with a syringe-based ExoFilter, the system is transformed into a new hybrid configuration by connecting two filters with distinct extraction mechanisms in a straightforward manner.

The key feature of this hybrid system is that bidirectional flow is provided by syringes at both ends. In this reciprocating flow environment, the strengths of each method are enhanced, creating an unprecedented synergistic effect. Specifically, in the ExoFilter, numerous small EVs in the sample are captured, reducing the burden on the TFF’s fine pores from clogging. In addition, as the liquid containing impurities is gradually removed through the TFF, a high concentration of target EVs remains in the sample, enabling the ExoFilter to more easily capture the residual EVs. This reciprocating flow process is repeated at least four to five times until all residual liquid is expelled, resulting in a dramatic increase in EV capture efficiency via the ExoFilter and significantly improved impurity removal performance by the TFF, thereby achieving a synergistic effect through this hybrid mechanism, as shown in Fig. 1(d). The key point is that only substances that satisfy both isolation mechanisms are isolated through the application of these two distinct operating principles, as illustrated in Figure 1(e).

The ExoTFF workflow involves three key steps: sample filtration, elution, and recovery, as depicted in Figure 1(f). First, the sample containing EVs is loaded into the ExoFilter syringe and passed through the membrane as it flows to the TFF, where anionic EVs are captured by the cationic mesh, and smaller proteins are removed. After repeating this process 3-4 times to ensure full filtration, an elution buffer with 1M NaCl is introduced, causing EVs to detach from the mesh. These EVs are retained within the TFF system due to their size and are subsequently concentrated and recovered in a smaller volume of PBS. A key advantage of the ExoTFF system is its ability to exchange buffers seamlessly, ensuring compatibility with various downstream applications. It also provides flexibility in concentration, enabling precise adjustments based on experimental requirements.

### Verification of EV Identity in Particles Isolated by ExoTFF

As shown in Fig. 2(a–b), EVs isolated using the ExoTFF method were visualized via scanning electron microscopy (SEM) and transmission electron microscopy (TEM). The images revealed that the EVs exhibited a spherical morphology with an average diameter of approximately 155 nm and a well-defined phospholipid bilayer. Nanoparticle tracking analysis (NTA) further characterized the size distribution of the isolated EVs, ranging from 50 to 450 nm, consistent with exosomes and large EVs (Fig. 2(c)).

**Figure 2.**
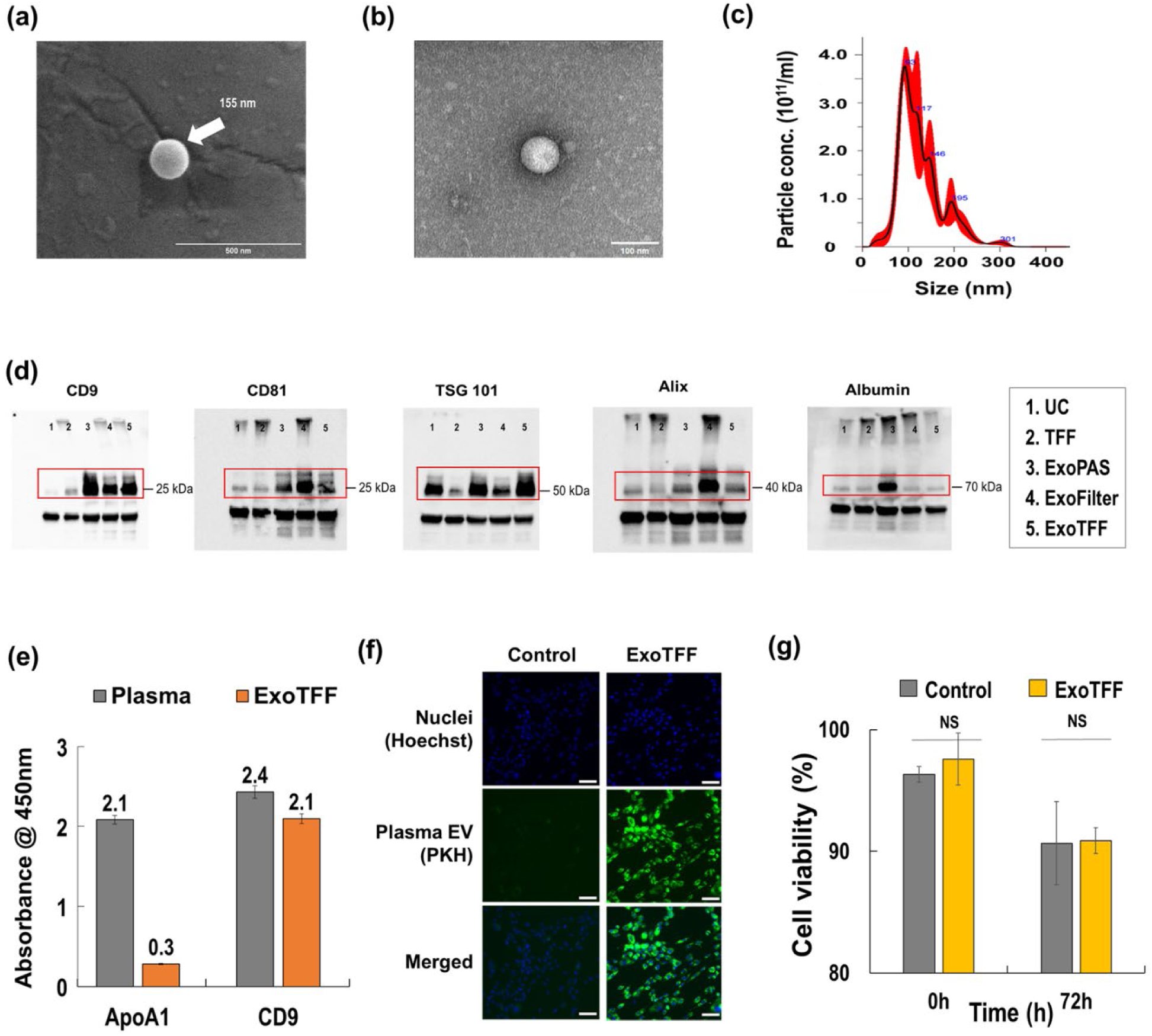
Characterization of EVs isolated from blood plasma using ExoTFF. (a) SEM image of an eluted EV, (b) TEM image of an eluted EV, (c) Particle size distribution and concentrations of EVs, (d) Western blot assay for isolated EVs including CD9, CD81, TSG101, Alix and Albumin, (e) ELISA Assay of ApoA1 as an HDL Marker and CD9 as an EV Surface Marker, (f) Ct values for miRNAs using RT-PCR, (g) Cellular uptake of EVs into human dermal fibroblasts (HDFs). (h) Cytotoxicity test of EVs in HDF after 72 hours.

To confirm the identity of the isolated EVs, Western blot analysis was performed to detect specific EV markers (Fig. 2(d)). The presence of CD9, CD81, TSG101, and Alix— well-established EV markers—was confirmed, while minimal albumin contamination indicated low plasma protein interference, demonstrating the high purity of EVs obtained using ExoTFF. Notably, albumin levels in ExoTFF isolates were comparable to those obtained via ultracentrifugation (UC), confirming that ExoTFF efficiently removes albumin without relying on UC.

Additionally, an ELISA assay validated the presence of CD9 as a positive biomarker of EVs, further confirming that the particles isolated by ExoTFF were indeed EVs rather than other types of nanoparticles. The assay also evaluated Apo-A1, a biomarker for high-density lipoprotein (HDL), which is typically abundant in plasma and challenging to eliminate with many conventional isolation techniques. As shown in Fig. 2(e), the Apo-A1 biomarker for HDL was significantly reduced to 14.3% of its original concentration in plasma after ExoTFF, despite its negative zeta potential. These findings in Fig. 2(d-e) represent a significant advancement, as many existing EV isolation methods struggle to sufficiently eliminate HDL and plasma proteins, further underscoring the high purity and effectiveness achieved with ExoTFF.

The functional bioactivity of EVs isolated by ExoTFF was confirmed through various assays, as depicted in Fig. 2(f). Cellular uptake studies demonstrated that human dermal fibroblasts effectively internalized the EVs. Additionally, cytotoxicity assays conducted over 72 hours (Fig. 2(g)) showed that the EVs did not induce significant cytotoxic effects, maintaining high cell viability.^18,19^ These results confirm that the isolated EVs retain their functional bioactivity without adversely affecting the cells.

**Figure 3.**
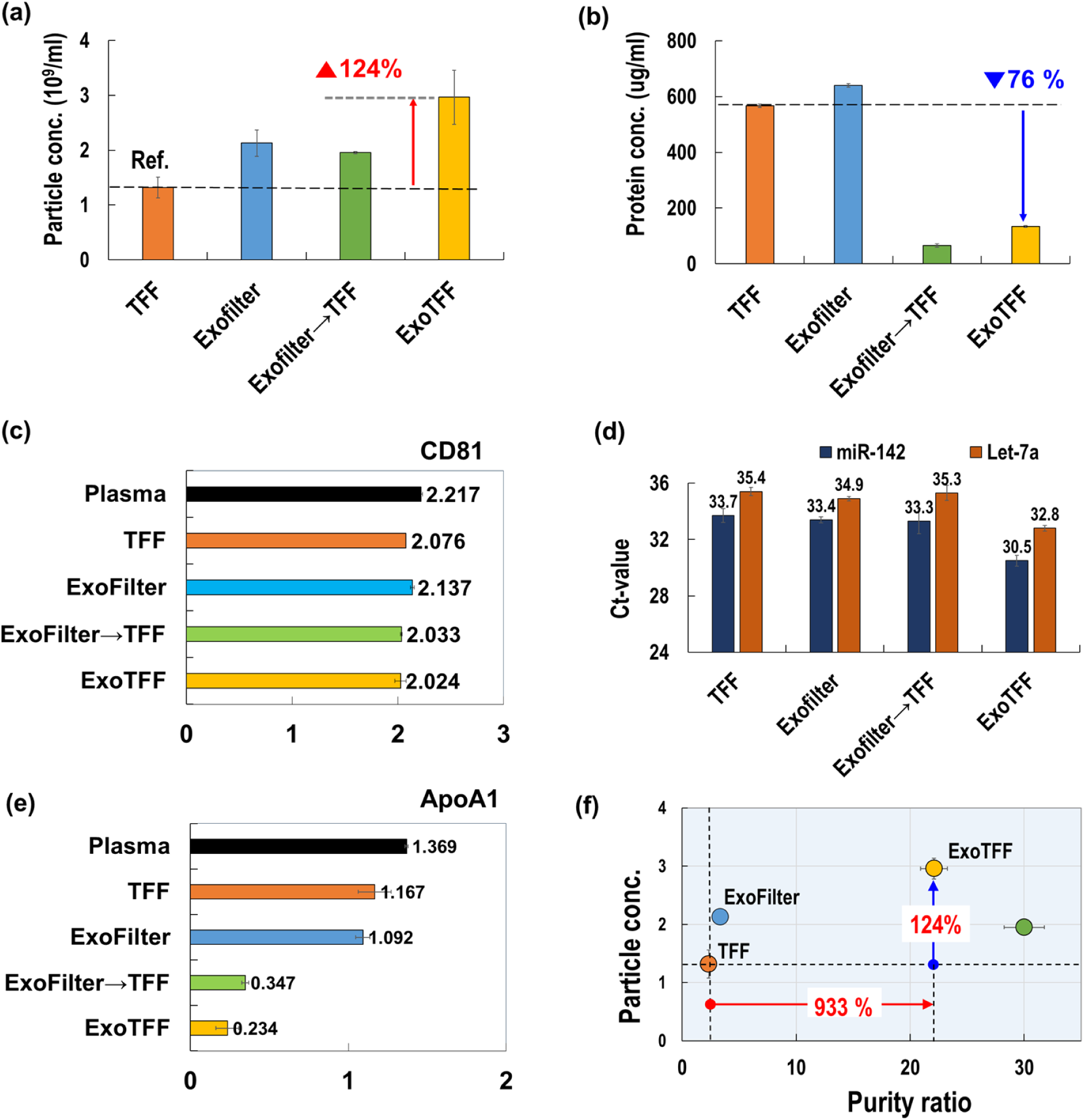
Comparative analysis of EV isolation methods (TFF, ExoFilter, ExoFilter→TFF, and ExoTFF) using plasma samples. (a) Particle concentration (b) Protein concentration of isolated EVs, (c) ELISA assays for specific EV markers (CD81), (d) Quantification of EV-derived miRNAs (miR-142 and let-7a) using RT-PCR, (e) ELISA assays for specific HDL markers (ApoA1), (f) Comparison of Purity ratios for various methods.

### ExoTFF Outperforms Other Methods in Yield and Purity

The superior effectiveness of ExoTFF was demonstrated through NTA and BCA analyses of plasma samples (Figures 3a–b), compared to various other methods. Compared to TFF as a reference, ExoTFF resulted in a remarkable 124% increase in particle concentration and a substantial 76% reduction in protein contamination.

The effectiveness of ExoTFF was further confirmed through ELISA analysis using three biomarkers: CD81 (an EV biomarker), ApoA1 (an HDL biomarker), and ApoB100 (a biomarker for VLDL and LDL). Plasma served as the reference, while TFF, ExoFilter, and ExoTFF were compared. Figure 3(c) presents CD81 expression levels as an indicator of EV yield. ExoTFF achieved a relative efficiency of 91.3% compared to plasma, while TFF and ExoFilter yielded similar results, suggesting comparable performance for EV isolation. However, unlike the NTA results shown in Figure 3(a), the ELISA data in Figure 3(c) did not reveal significant differences among the methods. This discrepancy may be attributed to inherent limitations in the properties of the biomarkers and the ELISA detection method—such as sensor surface saturation—which could restrict its ability to accurately reflect differences in extraction efficiency.

Therefore, in this study, we attempted a quantitative comparison of the extraction methods by amplifying miRNAs (miR-142 and let-7a), known as housekeeping genes within EVs, using RT-PCR. As shown in Figure 3(d), the Ct values for miR-142 and let-7a were significantly lower in samples processed by ExoTFF compared to TFF. For let-7a, the ΔCt value decreased by 2.6 between TFF and ExoTFF, indicating that ExoTFF isolates approximately six times more EVs than TFF.

Figure 3(e) shows the levels of ApoA1—an established HDL biomarker—measured by ELISA to assess HDL contamination. Initially, the HDL removal efficiency of TFF alone was only 15%, while that of ExoFilter alone was merely 20%. However, when the two techniques were sequentially combined in an “ExoFilter → TFF” configuration, the HDL removal efficiency dramatically increased to 75%. Moreover, employing the ExoTFF system with recirculation further boosted the HDL removal efficiency to 83%, demonstrating a remarkable synergistic effect.

When using TFF alone, the filtration process primarily relies on size. However, the actual hollow fiber membranes exhibit a wide distribution of pore sizes and their surfaces are predominantly negatively charged.^20^ Although the average size of HDL (8–12 nm) is theoretically much smaller than the 50 nm cutoff used in TFF—suggesting that HDL should easily pass through—in practice, a repulsive force between the negatively charged membrane surface and HDL may prevent effective removal through the TFF pores, causing these particles to remain in the retentate. Moreover, the actual operating conditions of the membrane and the heterogeneity in pore size distribution further contribute to the low HDL removal efficiency.

Figure 3(f) illustrates the relationship between particle concentration (vertical axis) and purity ratio (horizontal axis) for TFF, ExoFilter, and ExoTFF. The purity ratio is defined as the proportion of EVs relative to total protein or other contaminants, indicating how “clean” the isolated EVs are. The results show a substantial increase in both purity and yield with ExoTFF. While ExoFilter achieves a moderate improvement in particle concentration and purity ratio compared to TFF, ExoTFF demonstrates a dramatic rise: the particle concentration is 124% higher than that of TFF, and the purity ratio improves by 933%. Furthermore, ExoTFF’s hybrid process—combining selective capture and size-based filtration—yields a significantly higher purity ratio than TFF or ExoFilter alone, resulting in minimized protein contamination and maximized EV yield. Overall, Figure 3(f) underscores ExoTFF’s superior performance in delivering both high particle concentration and high purity ratio, confirming its effectiveness in producing cleaner, more concentrated EV preparations compared to TFF or ExoFilter alone.

One possible explanation for the remarkable synergistic HDL removal observed in the ExoTFF system is as follows: *1) Washing Process*: During the ExoFilter stage, negatively charged particles—including HDL—are captured. When connected to the TFF system, the subsequent washing step may preferentially remove HDL particles with relatively lower charge densities. These weakly bound particles can be dislodged and pass through the TFF pores, while strongly bound EVs remain retained. *2) Elution Process*: In the elution step, the high NaCl concentration in the buffer may cause Na⁺ ions to adsorb onto the TFF membrane pores. This adsorption can neutralize or screen the negative charges on the pore surfaces, effectively “opening” the pores to HDL. As a result, the repulsive interactions that would normally hinder HDL removal are reduced, allowing these particles to be efficiently cleared. Together, these mechanisms likely account for the enhanced HDL removal efficiency observed with the ExoTFF system compared to TFF or ExoFilter alone. However, further experimental validation is necessary to substantiate this hypothesis and fully elucidate the underlying mechanisms.

**Figure 4.**
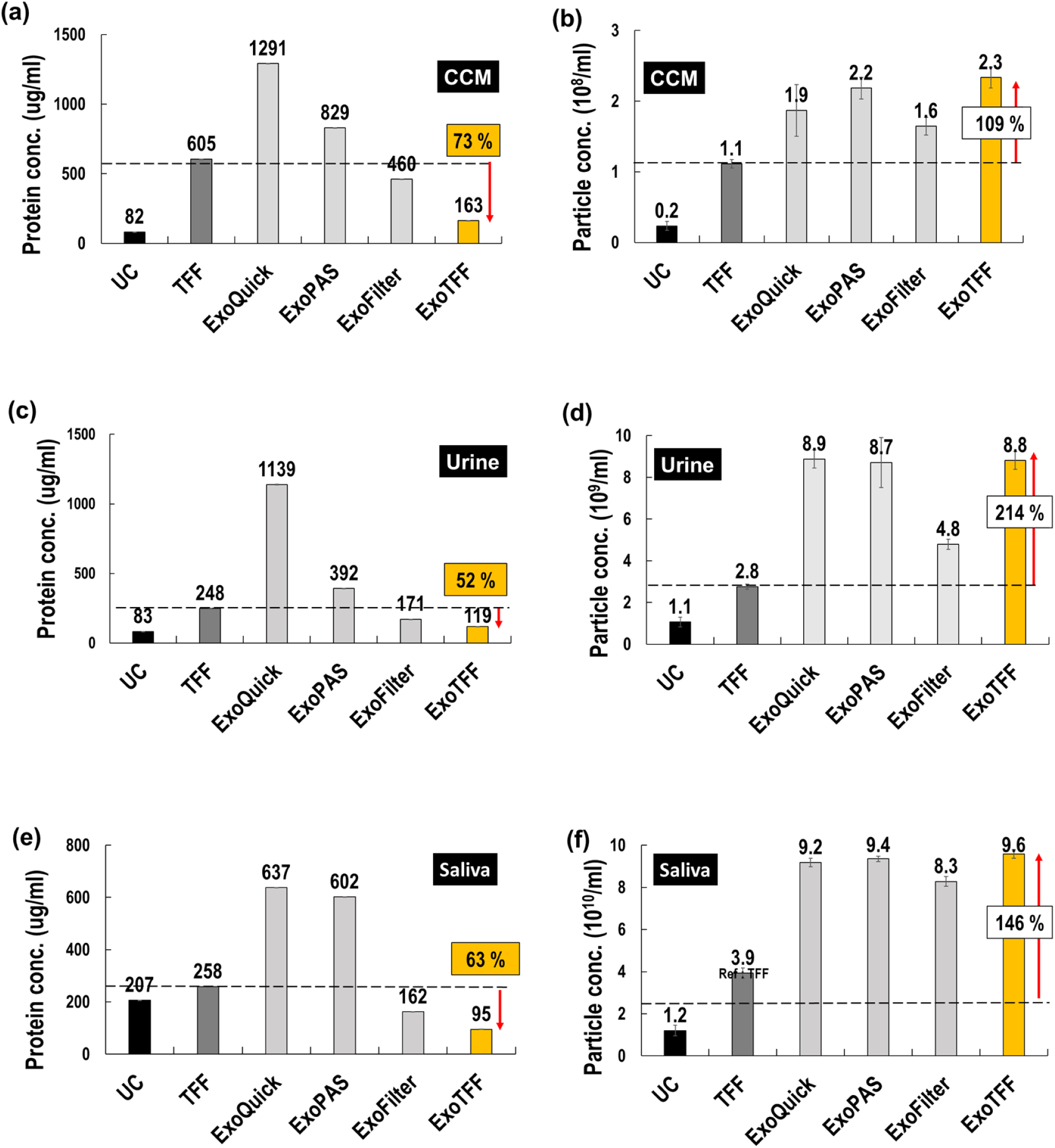
Comparison of EV Isolation Methods Across Various Biofluids (CCM, Urine, Saliva). Protein and particle concentrations of EVs isolated using different methods (UC, TFF, ExoQuick, ExoPAS, ExoFilter, and ExoTFF) from (a-b) cell-conditioned media (CCM), (c-d) urine, and (e-f) saliva samples.

### ExoTFF Outperforms Across Various Biofluids

We conducted further analyses to evaluate the ability of the ExoTFF method to isolate extracellular vesicles (EVs) from various biofluids, including conditioned cell media (CCM), urine, and saliva, and compared its performance against traditional methods. For CCM (Figures 4a-b), ExoTFF demonstrated a 73% reduction in protein concentration and a 109% increase in particle concentration compared to TFF, outperforming all other EV isolation methods tested, including UC, ExoQuick, ExoPAS, and ExoFilter.

In urine samples (Figures 4c-d), ExoTFF achieved a 52% reduction in protein concentration and a 214% increase in particle concentration compared to TFF, marking the highest performance among all methods including UC, ExoQuick, ExoPAS, and ExoFilter. For saliva samples (Figures 4e-f), ExoTFF resulted in a 63% decrease in protein concentration and a 146% increase in particle concentration relative to TFF, similarly showing superior performance over UC, ExoQuick, ExoPAS, and ExoFilter.

Overall, the results depicted in Figure 4 confirm that the ExoTFF method is highly effective at isolating EVs from a diverse range of biofluids, significantly enhancing both yield and purity compared to conventional isolation methods. The ExoTFF system not only demonstrates exceptional performance in blood plasma but also consistently outperforms conventional isolation methods in other biofluids, such as conditioned cell media (CCM), urine, and saliva. These findings indicate that ExoTFF is a versatile and powerful tool for EV isolation, offering broad potential utility in both research and clinical applications. Nevertheless, further experimental validation and mechanistic studies are warranted to fully elucidate the factors underlying its enhanced performance across various biofluids.

**Table 1.**
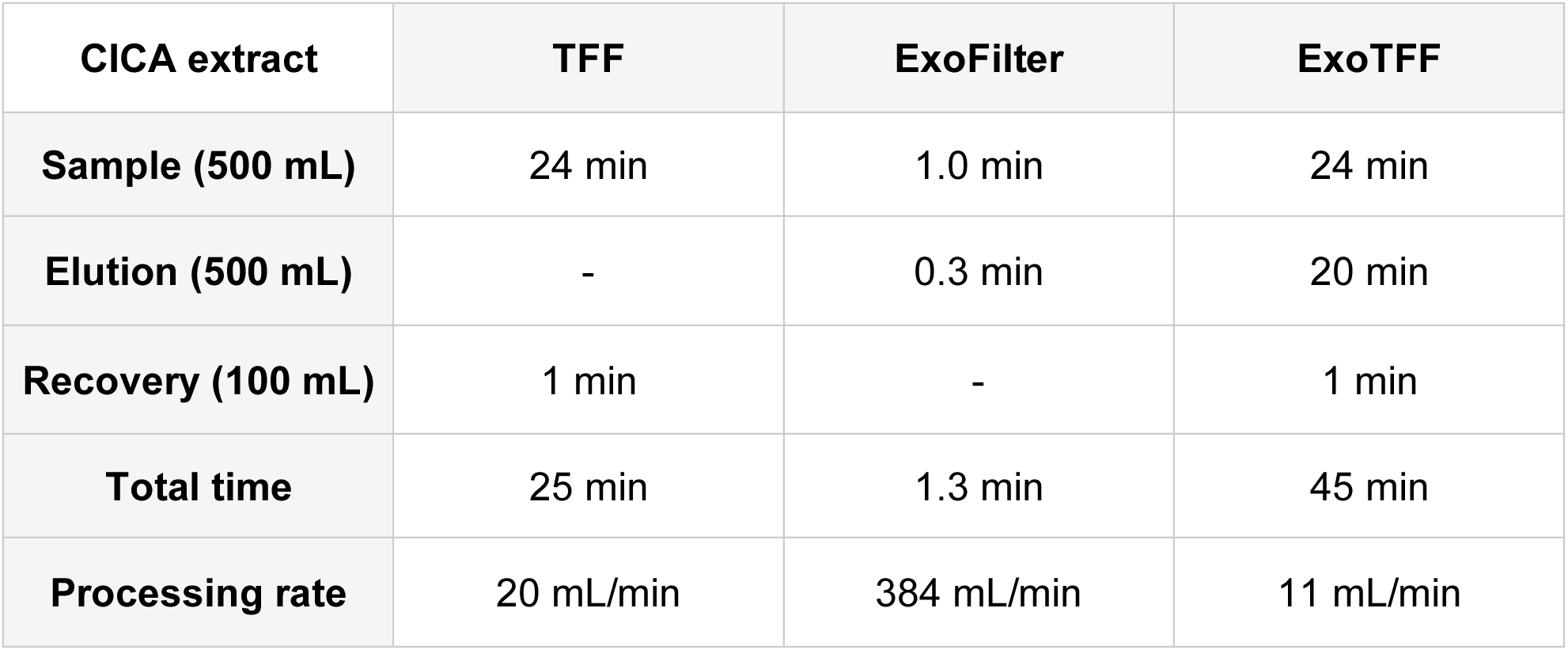
Operational thruput of three systems including TFF, ExoFilter and ExoTFF.

| CICA extract | TFF | ExoFilter | ExoTFF |
| --- | --- | --- | --- |
| Sample (500 mL) | 24 min | 1.0 min | 24 min |
| Elution (500 mL) | - | 0.3 min | 20 min |
| Recovery (100 mL) | 1 min | - | 1 min |
| Total time | 25 min | 1.3 min | 45 min |
| Processing rate | 20 mL/min | 384 mL/min | 11 mL/min |

### Automated ExoTFF System for Reliable, High-Throughput EV Isolation

In this study, the ExoTFF system was implemented as an automated setup consisting of a peristaltic pump, reservoir, ExoFilter and TFF and its performance was compared to that of automated TFF and ExoFilter systems, respectively. The sample volume used for the comparison was 500 mL of CICA (Centella Asiatica) plant extract. The performance comparison of the TFF, ExoFilter, and ExoTFF systems was analyzed, focusing on the operational characteristics of each system, including filtration speed, elution, recovery, and total processing time (Table 1). As a recent study reported, ExoFilter can process samples at an impressive rate of 384 mL/min using a charge-based filtration system. Additionally, the TFF system operates at a rate of 20 mL/min, focusing on size exclusion. Meanwhile, ExoTFF, which combines TFF and ExoFilter, requires a relatively longer time of 45 minutes to process 500 mL, resulting in a processing rate of 11 mL/min (or 660 mL/h).

The processing rate of the ExoTFF system represents a remarkable improvement compared to conventional methods. Moreover, as the ExoTFF system connects TFF and ExoFilter in series, increasing the capacity of each component can substantially enhance the system’s hourly throughput. This highlights the potential of ExoTFF to significantly improve high-purity EV processing in a high-throughput manner.

## 3. Discussion and conclusion

To fully understand the translational significance of extracellular vesicles (EVs) in health and disease, it is essential to develop reliable, scalable methods for isolating high-purity EVs. In this study, we introduce ExoTFF, a platform designed for high-throughput, scalable EV isolation with superior purity. Our results demonstrate high signal intensities for EV markers and miRNAs, with minimal contamination from non-EV proteins, underscoring the method’s effectiveness. By integrating two isolation principles—size and electrical charge—ExoTFF enables the selective purification of EV subpopulations that meet both criteria. In contrast, isolation technologies based on a single principle, such as size-exclusion chromatography, cannot achieve the same level of purity as dual-principle approaches. Moreover, ExoTFF’s combination of TFF and ExoFilter allows for the rapid processing of large sample volumes, providing high-throughput capacity for efficient, large-scale EV isolation from various biofluids.^16^ The system’s automated operation and fraction collection ensure consistent workflows and reproducible results across different sample types and volumes, making ExoTFF a robust solution that surpasses conventional EV extraction techniques.

In this study, ExoTFF demonstrated an impressive 933% improvement in purity over TFF, as shown in Fig. 3(f), showcasing the synergy between TFF and ExoFilter in isolating pure EVs. Remarkably, ExoTFF achieved higher purity ratios than both ultracentrifugation (UC) and TFF in complex biofluids such as conditioned culture media (CCM), urine, and saliva (Fig. 4). This enhanced performance stems from the charge-based filter, which selectively captures anionic particles, and size-based TFF, which effectively removes smaller proteins. As such, ExoTFF recorded the highest yields across all sample types. Given its proven efficacy, scaling up ExoTFF for industrial applications, particularly for processing volumes exceeding 200 liters, is a viable goal. The scalability of ExoFilter and the industrial applicability of large-volume TFF make this achievable.

However, the results also revealed discrepancies between quantification methods (NTA, BCA, RT-PCR, and ELISA), necessitating a reevaluation of the measurement techniques. For example, NTA includes particles ranging from 30 nm to 800 nm, which can count lipoproteins like LDL alongside EVs, potentially skewing the results. Similarly, BCA analysis quantifies total protein content, including both EV and contaminant proteins. Surprisingly, ELISA showed little difference in extraction yield, likely due to detection limits and potential interference from protein contaminants. On the other hand, RT-PCR, based on specific EV miRNA biomarkers, emerged as the most accurate quantification method, as RNase treatment ensured only internal miRNA from EVs was amplified. This highlights the importance of employing RT-PCR in future studies to obtain precise EV yield measurements.

Despite its advantages, ExoTFF faces challenges in isolating substances such as low-density lipoproteins (LDL) and very low-density lipoproteins (VLDL), which share similar sizes and charges with EVs. These lipoproteins coexist with EVs in biofluids and are difficult to separate using charge and size-based methods.^21^ Future research will focus on developing techniques to improve ExoTFF’s efficiency, including a new technology that targets and captures lipoprotein surface proteins. Success in this endeavor could revolutionize EV isolation by overcoming current limitations.

In summary, ExoTFF is an innovative platform for high-throughput, scalable EV isolation that integrates size and electrical charge to selectively purify EV subpopulations. These features offer the potential to accelerate EV research and expand the application of EVs in clinical and therapeutic fields.

## 4. Materials and Methods

### Preparation of cationic polymer coated mesh

Nylon meshes (Lixin Huarun MESH Co., China) were cut to 11 mm diameter using a laser cutter (BEAMO, MIRTECH Korea) and treated with protamine sulfate (Sigma-Aldrich, USA). First, the meshes were incubated in 0.1N hydrochloric acid for 30 minutes and washed with deionized water. They were then treated with 2.5% glutaraldehyde for 30 minutes, followed by additional rinsing. For activation, 5 mL of 0.1M EDC and NHS solutions were added, and the meshes were incubated for 1 hour before washing again. Finally, 100 mg of protamine sulfate dissolved in 10 mL of water was applied to the meshes, which were mixed for 15 minutes and left at room temperature for 6 hours. The coated meshes were rinsed three times to remove residual protamine sulfate.

### Design of the Syringe-Type ExoFilter

We developed a syringe-based ExoFilter system, consisting of multi-layered meshes conjugated with cationic substances, securely anchored within a 10 mL syringe using a frit. The capacity of the syringe can be adjusted to accommodate different sample volumes, enabling the production of various sizes of syringe-type ExoFilter. For this study, a 10 mL syringe was selected based on its compatibility with the minimum volume (5 mL) that can be processed by Tangential Flow Filtration (TFF), facilitating integration with the TFF system.

### Preparation of various samples

This study adhered to the ethical standards of the Declaration of Helsinki. Plasma samples were sourced from Zen-Bio Inc. (Research Triangle, NC, USA). Additional samples, including cell culture media from umbilical cord mesenchymal stem cells, urine, and saliva, were collected in sterile containers and purified through centrifugation. All samples were centrifuged at 3000 × g for 15 minutes to sediment and remove large particulates. The clarified supernatant was then passed through an 800 nm pore-sized mesh to exclude particles larger than 800 nm. Samples were subsequently stored at -80°C pending further analysis.

### EV Isolation via ExoFilter

The charge-based ExoFilter system was used to isolate EVs from plasma, urine, saliva, and CCM, with filters available in various capacities (1 mL, 3 mL, 15 mL, 200 mL) to suit different sample volumes. For smaller samples (1 mL, 3 mL, 15 mL), filtration was performed at 1 atm, followed by centrifugation at 5,000 × g for 1 minute, capturing negatively charged EVs through electrostatic interaction with the positively charged mesh. Elution was achieved with 1 M NaCl, using 1/5 of the sample volume. For larger 200 mL samples, the flow rate was controlled with a vacuum pump at - 68 kPa, followed by the same elution process. Protocol and operating principle of the syringe-based filtration system were depicted in Figure S2. The particle concentration increased progressively with each repeated process.

### EV Isolation via Tangential Flow Filtration (TFF)

For size-based filtration via tangential flow filtration (TFF), we utilized TFF-EVs (Hansa BioMed, Estonia) with an 800 kDa molecular weight cut-off and a fiber pore size of 50 ± 10 nm. We loaded 10 mL of various biofluids into the TFF system for processing. The samples were circulated through the TFF device using a syringe, repeatedly passing them back and forth until all the liquid was filtered. Following the initial filtration, a washing step involved circulating 10 mL of PBS in the same manner to cleanse the system. Finally, EV elution was performed by passing back and forth 2 mL of PBS through the TFF device back and forth to ensure comprehensive elution of the EVs.

### EV Isolation via ExoTFF

The ExoTFF system was used to isolate EVs from various samples, including plasma, urine, saliva, and cell culture media (CCM). During filtration, a 10 mL sample is loaded into the syringe-type ExoFilter and connected to the TFF system, with repetitive piston movements capturing negatively charged EVs in the positively charged mesh. Impurities like lipoproteins and proteins smaller than 30 nm are removed via TFF. For elution, 10 mL of 1M NaCl releases the captured EVs, with the process repeated until all liquid is expelled. In the recovery step, a new syringe with 2 mL of PBS recovers the EVs, concentrating them approximately fivefold.

### EV isolation via Ultracentrifugation (UC)

Ultracentrifugation (UC), the gold standard for EV isolation, is time-intensive, taking over 6 hours. For this process, 1 mL of biofluids was diluted 1:3 in PBS. The samples were centrifuged at 3,000 × g for 15 minutes, followed by 12,000 × g for 30 minutes to remove larger vesicles and debris. The supernatant was then ultracentrifuged at 120,000 × g at 4°C for 2 hours. After discarding the supernatant, the pellet was washed in PBS and spun again at 120,000 × g for 1 hour. The final pellet was resuspended in 200 µL of PBS for further analysis.

### EV isolation via ExoQuick

For polyethylene glycol (PEG)-based extracellular vesicle (EV) isolation, the ExoQuick exosome precipitation solution (EXOQ5A-1; System Biosciences, Palo Alto, CA, USA) was used. A total of 252 µL of the ExoQuick solution was added to 1 mL of biofluid sample and incubated for 30 minutes at 4°C. After incubation, the sample was centrifuged at 1500 × g for 30 minutes, and the supernatant was carefully decanted, leaving the pellet in the tube. To ensure complete removal of the ExoQuick solution, an additional centrifugation at 1500 × g for 5 minutes was conducted. The pellet was then resuspended in 200 µL of PBS.

### EV isolation via ExoPAS

The ExoPAS (Microgentas Inc., Seoul, Korea) isolates EVs from liquid samples utilizing a sequential application of protamine sulfate and polyethylene glycol (PEG). Initially, 1 mg of protamine sulfate is dissolved in 1 mL of deionized water and subsequently added to 1 mL of the biofluid sample. This process results in negatively charged EVs clustering, which is then enhanced by the addition of PEG 8000 (Sigma-Aldrich, USA). PEG 8000 solution (50%, 250 µL) is used to achieve a final concentration of 5.6%. The mixture is incubated at 4°C for 30 minutes. Following incubation, the sample is centrifuged at 3000 × g for 30 minutes. The supernatant is then carefully removed with a pipette, and the resultant EV pellet is resuspended in 200 µL of either PBS or 1 M NaCl solution for further analysis.

### Scanning Electron Microscopy (SEM) Imaging

SEM imaging was performed using an anodic aluminum oxide (AAO) membrane to analyze EVs isolated via ExoTFF from plasma. EVs were filtered through the membrane, fixed with glutaraldehyde for 30 minutes, and dehydrated stepwise with ethanol (25–100%). After oven-drying at 37°C for 2 hours, the membrane was platinum-coated to enhance contrast. EVs and aggregates were examined using a Quanta 250 FEG SEM (FEI, USA).

### Transmission Electron Microscopy (TEM) Imaging

The Carbon Formvar Film-150 copper grid was handled with tweezers to ensure the sample side faced up and placed on a Petri dish. EVs isolated by ExoTFF from plasma were applied (15 µL) and left to adsorb for one minute, covered to prevent contamination. For negative staining, 1% uranyl acetate was applied vertically, and excess stain was blotted off. After air drying on filter paper, the grid was imaged using a JEM-1400 Flash TEM (JEOL Ltd., Japan) at 120 kV.

### Nanoparticle Tracking Analysis (NTA)

Nanoparticle Tracking Analysis (NTA) was performed using the NS300 system with NTA 3.4 Software (NanoSight, UK) to assess EVs. Samples were diluted in pre-filtered PBS, and three 30-second videos were recorded per sample with the camera level at 14 and detection threshold at 11. This analysis determined the EVs’ average size and concentration based on the applied dilution factors.

### Bicinchoninic Acid assay (BCA)

Protein concentrations were measured using the Pierce™ BCA Protein Assay Kit (Thermo Scientific, USA) with a calibration curve prepared from bovine serum albumin (0–2000 μg/mL). Each standard and sample was measured in triplicate. A 100 μL sample was mixed with 2.0 mL of assay reagent, incubated at 37°C for 30 minutes, and allowed to cool to room temperature before absorbance was measured at 562 nm using a DS-11 spectrometer (Denovix, USA). Protein concentrations were calculated by comparing sample absorbance to the standard curve.

### Western Blotting Analysis

Proteins for Western blot analysis were extracted from EVs suspended in 200 µL of elution buffer, mixed with Laemmli buffer and 2-mercaptoethanol, and heated at 95°C for 10 minutes. SDS-PAGE was performed using a Mini-PROTEAN® TGX™ Precast Gel (Bio-Rad, USA). Western blotting employed antibodies against EV markers CD9, CD81, TSG101, ALIX, and albumin as a negative control. Antibodies included anti-CD9, anti-CD81, anti-TSG101, and anti-ALIX (Abcam, UK), with goat anti-rabbit IgG as the secondary antibody. Protein bands were visualized using the ChemiDoc™ XRS+ System with enhanced chemiluminescence (ECL) detection.

### Immunocapture-based ELISA

Anti-CD9 and anti-CD81 antibodies (R&D Systems, USA) were diluted to 5 µg/mL in PBS, with 100 µL added to microwells and incubated at 37°C for 2 hours. After washing, 0.5% casein was added and incubated for 1 hour. Exosome samples, diluted in 0.5% casein and 0.1% Tween-20 to various concentrations, were added to the wells and incubated overnight at 37°C. Following washing, biotinylated anti-CD63 antibody (BioLegend, USA) was applied for 2 hours, then streptavidin-HRP (Fitzgerald, USA) was added for 1 hour. The reaction was developed with HRP substrate for 13 minutes, stopped with sulfuric acid, and absorbance was measured at 450 nm using a SPECTROstar Nano microplate reader.

### miRNA Analysis via RT-qPCR

To quantify specific miRNA markers within EVs, RNA extracted from the EVs was reverse transcribed using the TaqMan MicroRNA RT kit (4366596, Life Technologies, Eugene, OR, USA) and analyzed with TaqMan MicroRNA Assays (4427975, Life Technologies, USA). The assay specifically targeted hsa-let-7a-5p and hsa-miR-142-3p, using TaqMan Universal Master Mix II without UNG (4440040, Life Technologies, Eugene, OR, USA).

### Cellular Uptake Analysis

Cellular uptake of EVs by human dermal fibroblasts (HDF) was assessed using EVs labeled with PKH67 green fluorescent dye (Sigma-Aldrich, Burlington, MA, USA). The labeling was conducted for 15 minutes at 25°C, followed by the removal of excess dye through a 100-kDa filter. The HDF cells were then exposed to PKH67-stained EVs at a concentration of 2 × 10^9^ particles/mL in their culture medium. Nuclear staining was achieved by adding Hoechst 33342 dye to the medium, and the cellular uptake of EVs was visualized using a fluorescence microscope (Eclipse Ti2; Nikon, Tokyo, Japan).

### Cytotoxicity test

To evaluate cell viability, HDF cells were seeded in 96-well plates at a density of 1 × 10⁴ cells/cm² and allowed to adhere for 24 hours. Following this, the cells were washed and then treated with EV-depleted FBS-supplemented medium along with EVs at a concentration of 2 × 10⁹ particles/mL for 72 hours. Cell viability was evaluated using a WST-1 assay kit (EZ-Cytox; DoGenBio, Seoul, Korea). The WST-1 reagent was mixed with the culture medium in a ratio of 1:10, and 100 µL of the mixed solution was added to each well of the 96-well plate. The cells were incubated at 37°C for 1 hour before the absorbance at 450 nm was measured to ascertain viability.

### Zeta potential analyzer

Zeta potentials for EVs and various plasma proteins including albumin, γ-globulin, and fibrinogen were determined using a Zetasizer Pro (Malvern Panalytical, Malvern, UK). Given the challenges associated with resuspending EVs and plasma proteins in deionized water, each 10-µL sample was diluted in 990 µL of deionized water prior to measurement. The cationic nylon mesh was analyzed using a Surpass 3 analyzer (Anton Paar GmbH, Austria), employing a protamine-conjugated mesh measuring 20 mm by 10 mm for this purpose.

## AUTHOR INFORMATION

### Corresponding Author

**Sehyun Shin** − School of Mechanical Engineering, Department of Micro-Nanosystem Technology, Korea University, Engineering Research Center for Biofluid Biopsy, Seoul, 02841, Republic of Korea; phone: +82 2 3290 3377; Fax: +82 2 928 5825; ORCID ID: 0000-0002-2611-5610

### Authors

**YongWoo Kim** − School of Mechanical Engineering, Engineering Research Center for Biofluid Biopsy, Seoul, 02841, Republic of Korea

**SoYoung Jeon** − Department of Micro-Nanosystem Technology, Korea University, Engineering Research Center for Biofluid Biopsy, Seoul, 02841, Republic of Korea

**Kangmin Lee** − School of Mechanical Engineering, Engineering Research Center for Biofluid Biopsy, Seoul, 02841, Republic of Korea

### Author Contributions

Y.W.K. ‘contributed formal analysis, investigation, methodology, writing—original draft; K.M.L contributed investigation, methodology; S.Y.J. contributed investigation; methodology; S. S. contributed conceptualization; formal analysis; methodology; supervision; visualization; writing—review & editing.

### Biographies

YongWoo Kim is currently enrolled in the MSc program in the Department of Mechanical Engineering at Korea University. Since 2023, he has been researching aptamer-based EV isolation and reversible release under the guidance of Prof. Sehyun Shin at Korea University.

SoYoung Jeon received her MSc degree in Bioinformatic Engineering from Korea University in 2024. She subsequently joined the PhD program under the guidance of Prof. Sehyun Shin at Korea University, where she is developing EV isolation technologies as well as EV-derived miRNA diagnostics.

Kangmin Lee received his MSc degree from the Department of Mechanical Engineering at Korea University in 2025. Since 2023, he has been studying the development of EV isolation and characterization as a Master’s student under the guidance of Prof. Sehyun Shin at Korea University.

Sehyun Shin is a Professor in the School of Mechanical Engineering at Korea University. He received his BS and MSc degrees from Seoul National University and his PhD from Drexel University. Currently, he serves as the Director of the Engineering Research Center for Biofluid Biopsy. His recent research areas include platelet function assays, global hemostasis and thrombosis, RBC rheology, EV isolation, and molecular diagnostics for liquid biopsy applications.

## Acknowledgements

This project was conducted with the support of the Alchemist Project of the Korea Evaluation Institute of Industrial Technology (KEIT 20018560/NTIS 1415184668) funded by the Ministry of Trade, Industry & Energy (MOTIE, Korea) and National Research Foundation of Korea (NRF) Grant funded by the Korean Government, MSIP (RS-2023-00207833). The funders had no role in study design, data collection and analysis, decision to publish or preparation of the manuscript.

## Data availability

The main data supporting the results of this study are available within the manuscript and supplementary information files. The raw data files are available for research purposes from the corresponding author upon reasonable request. Source data are provided with this paper.

