## Supplementary material for "Advanced Extracellular Vesicle Isolation: A Hybrid Electrokinetic-Tangential Flow Filtration Approach for Improved Yield, Purity, and Scalability"

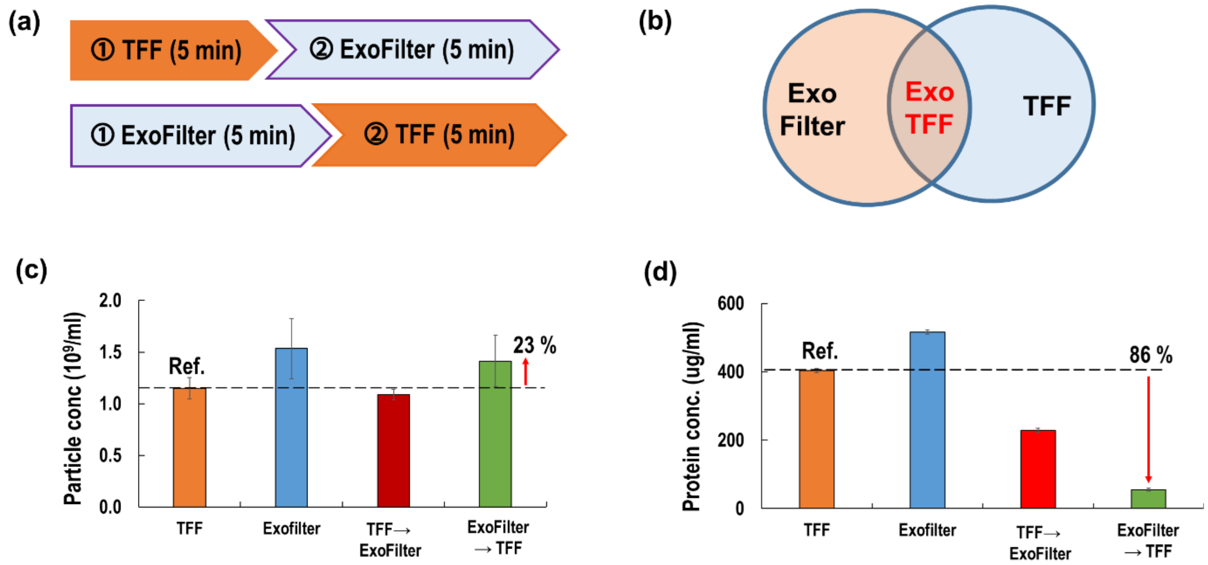


**Figure S1.** (a) Schematic representations of the order-dependent tests, comparing the sequence of TFF followed by ExoFilter and ExoFilter followed by TFF, (b) Application of dual principles in ExoTFF, (c-d) Results of the order-dependent tests, showing the effects on particle concentration and protein concentration from blood plasma.

In this study, while attempting to simply combine the two aforementioned technologies, we explored two different order-dependent tests: Comparison of TFF Followed by ExoFilter and ExoFilter Followed by TFF, as shown in Figures S1(a). This combined isolation method is expected to yield exceptionally pure EVs by applying two principles. The order of these techniques had little effect on the yield of EV isolation, but it had a significant impact on impurity removal. For instance, in the case of blood plasma (10 mL), treating the sample first with the ExoFilter and then with TFF led to a substantial reduction in protein concentration, achieving an 86% decrease compared to isolating EVs via TFF alone, as shown in Fig. S1(d). This result warrants more careful interpretation, as the underlying mechanism may involve multiple factors.

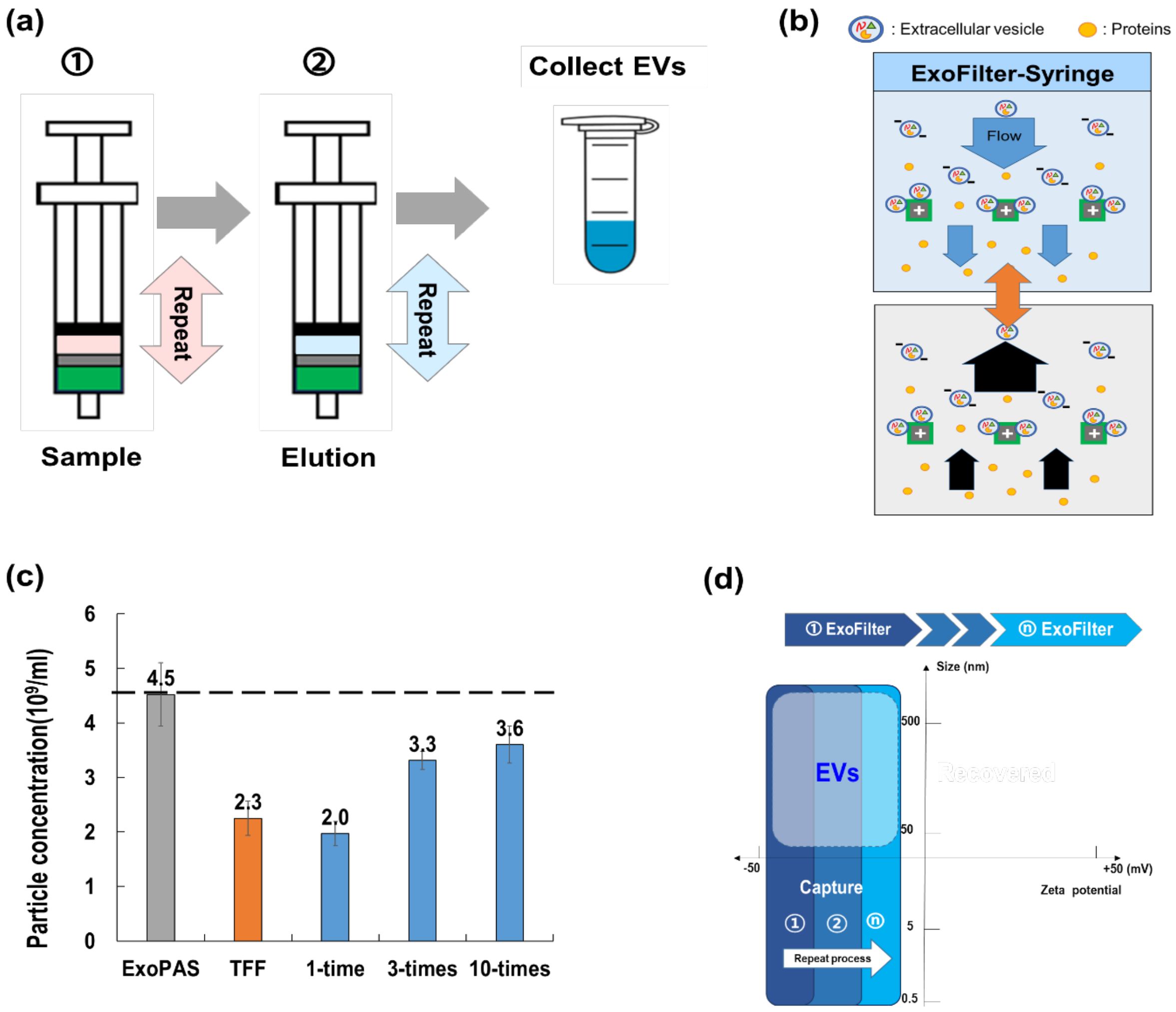


**Figure S2. Protocol and operating principle of the syringe-based filtration system.** (a) Workflow of the syringe-based filtration process, (b) Mechanism of particle capture in the syringe-based system, (c) NTA measurement showing increased particle concentration with repeated filtration cycles, (d) Effective EV isolation area within the filtration system.
